# Hepatitis C virus NS5A inhibitor daclatasvir allosterically impairs NS4B-involved protein-protein interactions within the viral replicase and disrupts the replicase quaternary structure in a replicase assembly surrogate system

**DOI:** 10.1101/335273

**Authors:** Yang Zhang, Jingyi Zou, Xiaomin Zhao, Zhenghong Yuan, Zhigang Yi

**Affiliations:** Key Laboratory of Medical Molecular Virology and Department of Medical Microbiology, School of Basic Medical Sciences, Shanghai Medical College, Fudan University, Shanghai, China; Department of pathogen diagnosis and biosafety, Shanghai public health clinical center, Fudan University, Shanghai, China

**Keywords:** Hepatitis C Virus: NS5A, NS4B, daclatasvir, direct-acting antivirals, replicase, quaternary structure, protein-protein interaction

## Abstract

Daclatasvir (DCV) is a highly potent direct-acting antiviral that targets the non-structural protein 5A (NS5A) of hepatitis C virus (HCV) and has achieved great clinical successes. Previous studies demonstrate its impact on the viral replication complex assembly. However the precise mechanism by which DCV impairs the replication complex assembly remains elusive. In this study, by using HCV subgenomic replicons and a viral replicase assembly surrogate system that expresses the HCV NS3-5B polyprotein to mimic the viral replicase assembly, we dissected the impacts of DCV on aggregation and tertiary structure of NS5A, the protein-protein interactions within the viral replicase and the quaternary structure of the viral replicase. We found that DCV didn’t affect aggregation and tertiary structure of NS5A. DCV induced a quaternary structural change of the viral replicase, evidenced by selectively increasing of the NS4B’s sensitivity to proteinase K digestion. Mechanically, DCV impaired the NS4B-involved protein-protein interactions within the viral replicase. The DCV-resistant mutant Y93H was refractory to the DCV-induced reduction of the NS4B-invoved protein interactions and the quaternary structural change of the viral replicase. In addition, Y93H reduced NS4B-involed protein-protein interactions within the viral replicase and attenuated viral replication. We propose that DCV may induce a position change of NS5A, which allosterically affects the protein interactions within the replicase components and disrupts the replicase assembly.

**Importance:** The development of the direct-acting antivirals (DAA) has resulted in great clinical achievements for Hepatitis C Virus (HCV) treatment. Daclatasvir (DCV) is an inhibitor targeting the non-enzymatic NS5A, with the 50% effective concentration values in the picomolar range. Accumulated data suggest that DCV blocks the biogenesis of the HCV replication complex. However the mechanistic actions of DCV are still largely unknown. Insights into the action mechanism of DCV on the viral replication complex assembly of HCV may enlighten the development of next generation of DAAs and new anti-viral strategies for other positive-strand RNA viruses for which there are a scarcity of DAAs. Herein, using HCV subgenomic replicons and a viral replicase assembly surrogate system, we dissected the mechanistic actions of DCV on the viral replicase assembly. We found that DCV allosterically impairs NS4B-involved protein-protein interactions within the viral replicase and disrupts the quaternary structure of the viral replicase.

## Introduction

Hepatitis C Virus (HCV) chronically infects approximately 160 million people worldwide and causes hepatocellular carcinoma (HCC) (1). HCV is a member of the *Flaviviridae* family. Its 9.6-kb positive-sense RNA genome encodes a single polyprotein, which is co- and post-translationally cleaved into at least 10 individual proteins. The protein order of the individual structural and non-structural (NS) proteins is: 5′-C-E1-E2-p7-NS2-NS3-NS4A-NS4B-NS5A-NS5B-3′ (see review, (2)). The non-structural protein 3 (NS3) has a N-terminal serine protease domain and a C-terminal RNA helicase/NTPase domain. The NS4A is the cofactor for the NS3 protease. The NS4B is a multi-spanning integral membrane protein. The NS5A is a multifunctional viral protein. The NS5B is an RNA-dependent-RNA polymerase (see review, (3)). The HCV non-structural viral protein NS3, NS4A, NS4B, NS5A and NS5B assemble the viral replicase in the endoplasmic reticulum (ER), which induces protrusion of the ER to form a double membrane vesicle (DMV) structure, namely the viral replication complex (4). Overexpression of the HCV polyprotein encompassing NS3 to NS5B induces the formation of the DMV structure, mimicking the replicase assembly seen in virally infected cells (4). Expression of NS5A alone, albeit less efficient, can also induce DMVs (4), suggesting a central role of NS5A in the replication complex assembly.

The development of the direct-acting antivirals (DAA) against the replicase components has resulted in great achievements for HCV treatment (5). Daclatasvir (DCV) is one of the inhibitors targeting NS5A, with the 50% effective concentration values in the picomolar range against HCV replicons from various genotypes (6). NS5A contains three domains. The N-terminal amphipathic helix (AH) mediates its membrane association (7). The domain I (DI) following the AH is highly structured and proposed to mediate NS5A dimerization and RNA-binding (8, 9). The domain II (DII) is intrinsically disordered and is involved in genome replication (10, 11). The domain III (DIII) is natively unfolded and is required for virion production (12, 13). DCV is likely targeting the DI of NS5A, as the DCV-resistant mutation Y93H resides in the DI (6, 14).

DCV was identified as an extremely potent inhibitor of HCV replication in cell-based replication assays (6). DCV or DCV-like inhibitors affect the hyper-phosphorylation of NS5A (15, 16) and alter the subcellular localization of the NS5A (15, 17, 18). By using an HCV replicase surrogate system, Berger *et al.* demonstrated that DCV blocks the early biogenesis of the viral membranous replication factories (19). In addition, DCV blocks transfer of viral genome to the assembly sites to affect virion assembly (20).

Assembling a viral replicase on modified host intracellular membranes to form a replication complex (RC) is a common strategy for viral replication of almost all of the positive-strand RNA viruses (21). Insights into the action mechanism of DCV on the viral replication complex assembly of HCV may enlighten the development of new anti-viral strategies for other positive-strand RNA viruses for which there are a scarcity of DAAs. In this study, we used HCV subgenomic replicons and a viral replicase assembly surrogate system wherein the HCV NS3-5B polyprotein is expressed to mimic the induction of the viral replicase (4). We dissected the mechanistic actions of DCV on the viral replicase assembly.

## Results

### DCV blocks HCV replication and reduces NS5A hyper-phosphorylation in replicon cells and in a replicase assembly surrogate system

To set the dosages of DCV used in the experiments, we first monitored HCV infection in the presence of various concentrations of DCV. We infected Huh7.5 cells with Jc1G, a genotype 2a virus expressing a secreted *Gaussia* luciferase (Gluc) reporter (22). DCV treatment at 10 nM nearly completely blocked viral infection (Fig. 1A and B). In a genotype 2a subgenomic replicon (sgJFH1) (22) cell line and a genotype 1b subgenomic replicon (BB7) (23) cell line, DCV treatment also reduced viral replication (Fig. 1C and D). As reported (15, 19), in the sgJFH1 replicon cells, DCV reduced hyper-phosphorylation of NS5A (Fig. 1D). DCV treatment for 4 hours reduced the hyper-phosphorylated form of NS5A without affecting the protein levels of NS3 and the basal-phosphorylation of NS5A (Fig. 1E), suggesting the reduction of NS5A hyper-phosphorylation is an early event of DCV action and maybe used as an indicator of the action of DCV. DCV treatment with levels greater than 10 nM didn’t further reduce the NS5A hyper-phosphorylation (Fig. 1E and F).

**Figure 1.**
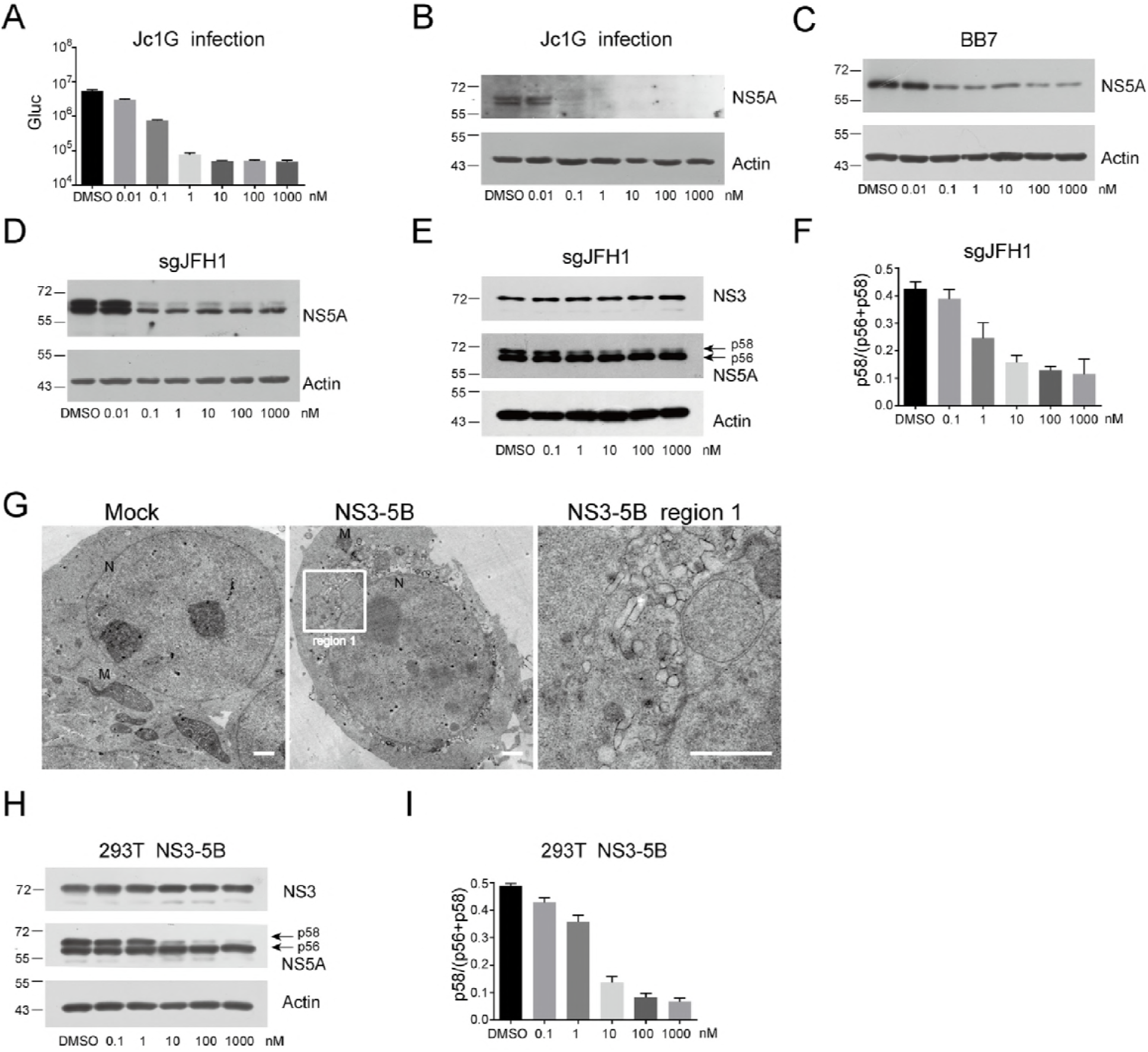
DCV blocks HCV replication and reduces NS5A hyper-phosphorylation. (A, B) Huh7.5 cells were infected with Jc1G at an MOI of 0.1 for 8 hours. After three washes with PBS, mediums with various concentrations of DCV were added. Forty-eight hours later, (A) the Gaussia luciferase (Gluc) activity in the supernatants was measured. Mean values ± SD are shown (n=3). (B) The cell lysates were analyzed by Western blotting with the antibodies indicated. (C) BB7 replicon cells and (D) sgJFH1 replicon cells were treated with various concentrations of DCV. After 48 hours, the cells were harvested for Western blotting analysis with the antibodies indicated. (E, F) Replicon cells (sgJFH1) were treated with various concentrations of DCV. (E) Four hours later, the cells were harvested for Western blotting analysis with antibodies indicated. (F) The hyper-phosphorylated form (p58) and the basal-phosphorylated form (p56) of NS5A were quantified. The ratio of the hyper-phosphorylated form of NS5A (p58) to the total NS5A (p56+p58) was calculated. Mean values ± SD are shown (n=3). (G) HEK293T cells were mock-transfected (Mock) or transfected with HCV NS3-5B expressing plasmid. After 24 hours, the cells were fixed, embedded and the ultrathin sections were prepared as described in material and methods. The samples were observed by transmission electron microscope. M, mitochondria; N, nucleus. Representative pictures are shown. Scar bar, 1μm. (H, I) HEK293T cells were transfected with plasmids expressing HCV NS3-5B in the presence of various concentrations of DCV. (H) Twenty-four hours later, the cells were harvested for Western blotting analysis. (I) The hyper-phosphorylated form (p58) and the basal-phosphorylated form (p56) of NS5A were quantified. The ratio of the hyper-phosphorylated form of NS5A (p58) to the total NS5A (p56+p58) was calculated. Mean values ± SD are shown (n=3). Representative data from multiple experiments with similar results are shown. The values to the left of the blots in panels B, C, D, E, H are molecular sizes in kilodaltons.

DCV blocks biogenesis of the HCV replication complex in a replicase assembly surrogate system (19). We took a similar strategy and expressed the HCV NS3-5B (JFH1) polyprotein in HEK293T cells. We selected HEK293T cell for certain reasons. First, it supports replication of the JFH1 subgenomic replicon (24), making it physiologically relevant to study the replicase assembly in these cells. Second, the expression of HCV NS3-5B polyprotein in HEK293T cells induces membranous web structures in the perinuclear regions (Fig. 1G), similarly as in Huh7 cells (4). Third, the high transfection efficiency in these cells facilitates the expression of the viral polyprotein to a similar level as found in the replicon cells (Data not shown and data described below). We then expressed HCV (JFH1) polyprotein encompassing the NS3 to NS5B (3-5B) (22) in HEK293T cells and treated the cells with DCV. As in the replicon cells, DCV reduced the hyper-phosphorylation of NS5A (Fig. 1H) and didn’t further reduce the NS5A hyper-phosphorylation above 10 nM (Fig. 1H and I). As DCV treatment didn’t further reduce the NS5A hyper-phosphorylation above 10 nM, we selected 10 nM for DCV treatment in the subsequent experiments.

### DCV does not trigger aberrant aggregation of NS5A

It has been reported that DCV treatment either change the size of the NS5A-clustering sites (15) or not (19). We assessed the impact of DCV on the NS5A distribution in either JFH1 subgenomic replicon cells or in HEK293T cells expressing con1b NS3-5B and found no effect of DCV on the sizes of the NS5A-punctae (Data not shown). We noticed that NS5A was redistributed to lipid droplet in the cells expressing the NS3-5B of genotype 2a (JFH1) (Data not shown), which is in consistent with a previous study (17).

Purified NS5A forms a dimer *in vitro* and there are studies implying that NS5As exist as polymers within the cells (8, 9, 25). It has been proposed that DCV may regulate the polymers of NS5A to block the replicase assembly. We recently identified VCP, a member of the ATPases associated with diverse cellular activities (AAA+ ATPase family), as an indispensable host factor for HCV replication (22, 26). Pharmacological inhibition of VCP resulted in aberrant aggregation of HCV NS5A and reduction of NS5A hyper-phosphorylation (26). As DCV also decreased NS5A hyper-phosphorylation, we investigated if DCV acts in a similar mechanism as VCP inhibitors. We treated the subgenomic sgJFH1 replicon cells with DCV or a VCP inhibitor NMS-873 (27), and then analyzed NS5A aggregation by semi-denaturing detergent agarose gel electrophoresis (SDD-AGE) as in our previous studies (26). The SDS-resistant protein aggregations appear as smears in the agarose gel due to their heterogeneity (28). NMS-873 treatment resulted in pronounced smears as reported (26) (Fig. 2A). In contrast, comparing with mock treatment (DMSO), DCV treatment didn’t significantly increase the aggregation of NS5A (Fig. 2A to 2C). Therefore, DCV reduces the hyper-phosphorylation of NS5A probably through a different mechanism other than affecting the aggregation of NS5A.

**Figure 2.**
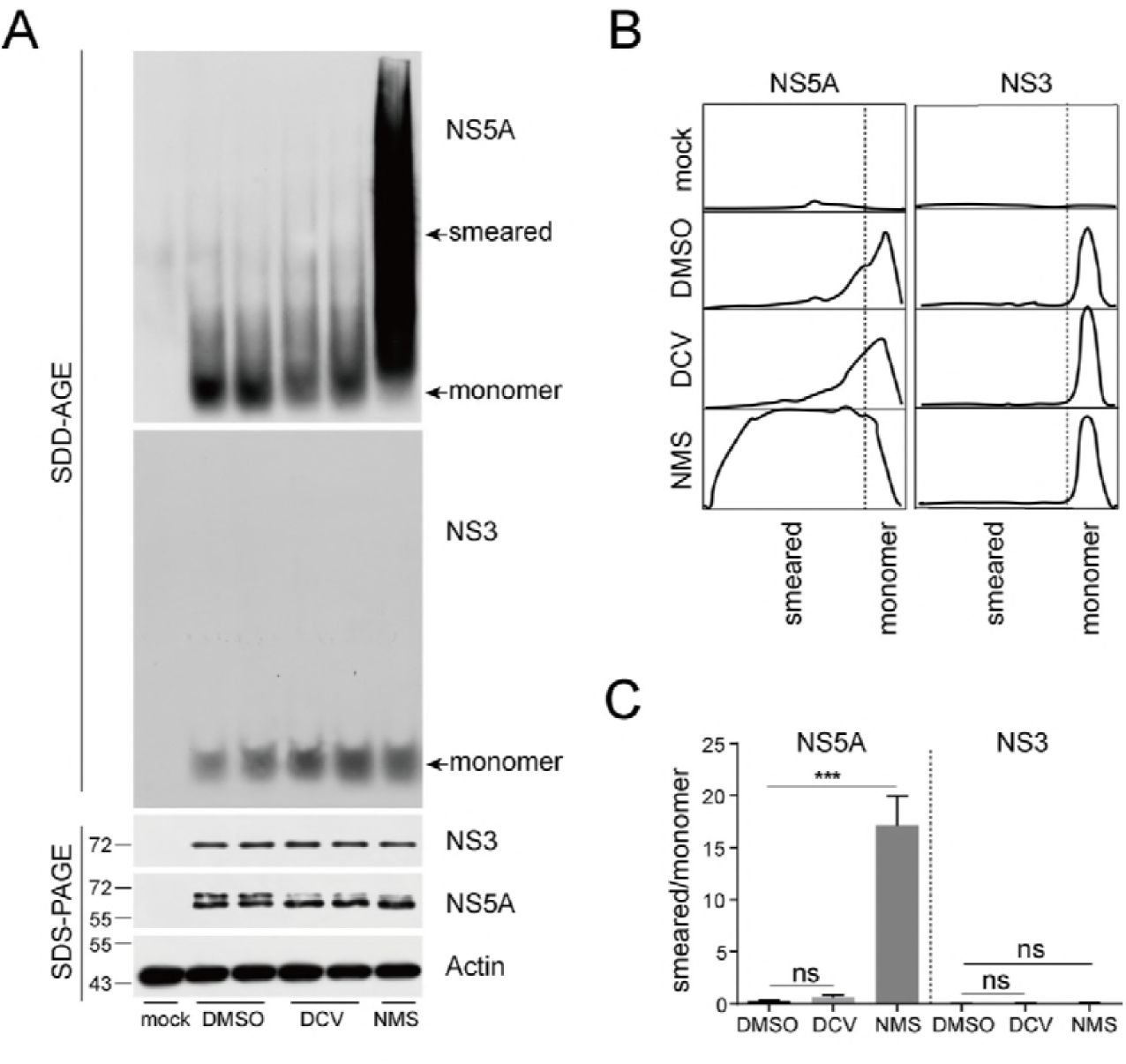
DCV does not trigger aberrant aggregation of NS5A. (A) The sgJFH1 replicon cells were treated with DCV (10 nM) or NMS-873 (5 μM) for 4h and then harvested for SDS-PAGE or SDD-AGE. In the SDD-AGE blot, the high-molecular-weight smears are labeled as ‘smearing’ and the low-molecular-weight monomers are labeled as ‘monomer’. (B) The relative density profile of each lane in the SDD-AGE blot from (A) was generated by Image J software. The peaks corresponding to the ‘monomer’ bands and the ‘smeared’ species were identified. Representative data from one lane from the duplicated samples are shown. (C) The density of the areas corresponding to the ‘monomer’ bands and the ‘smeared’ species were quantified and the ratio of the ‘smeared’ species to the ‘monomer’ bands (smeared/monomer) was calculated and plotted. Mean values ± SEM are shown (n=4). Statistical analysis was performed between the treated groups and the mock-treated groups (DMSO) as indicated. (ns, not significant; ***, *P*<0.001; two-tailed, unpaired *t*-test.). Representative data from multiple experiments with similar results are shown. The values to the left of the blots in panels A are molecular sizes in kilodaltons.

### DCV does not change the tertiary structure of NS5A protein

We then investigated the impact of DCV on the tertiary structure of NS5A. We monitored the structural changes of NS5A by limited trypsin proteolysis, a technique previously used to monitor the sgRNA binding-induced conformational change of CRISPR-Cas9 (29). The protein’s sensitivity to the enzyme digestion relies on the exposure of the cleavage sites and any changes in the proteolysis pattern reflect conformational changes in the tertiary structure of the protein. We treated the sgJFH1 replicon cells with DCV (10 nM) or the VCP inhibitor NMS-873 (5 μM) for 4 hours and then solubilized the cells using buffer containing 1% Triton X-100. The solubilized cell lysates were incubated with trypsin at various mass ratios. The digested cell lysates were separated by SDS-PAGE and the viral proteins were detected by Western blotting with an anti-NS5A monoclonal antibody and an anti-NS3 monoclonal antibody. NS5A from the DMSO-treated cells begun to demonstrate sensitivity to trypsin digestion when the trypsin was used at a mass ratio of 1:3000, as evidenced by the reduction of NS5A protein levels compared with the mock-treated groups (Fig. 3A). NMS-873 treatment increased NS5A’s sensitivity to trypsin, resulting in more reduction of the NS5A and at a lower mass ratio, compared with DMSO treatment (Fig. 3A). In contrast, NS3 and Actin were relatively resistant to trypsin digestion using these conditions (Fig. 3A), which is presumably due to the lower accessibility to trypsin. We quantified the proteolysis of NS5A and NS3 when digested the DCV- and NMS-873-treated cell lysates with trypsin at the mass ratio of 1:3000 (trypsin/total cellular protein). Comparing with DMSO treatment, DCV treatment only reduced the hyper-phosphorylation of NS5A but did not significantly affect NS5A’s sensitivity to trypsin (Fig. 3B and 3C). In contrast, NMS-873 treatment significantly increased NS5A’s but not NS3’s sensitivity to trypsin (Fig. 3B and 3C). These data indicate that DCV does not likely cause a change in the overall tertiary structure of NS5A.

**Figure 3.**
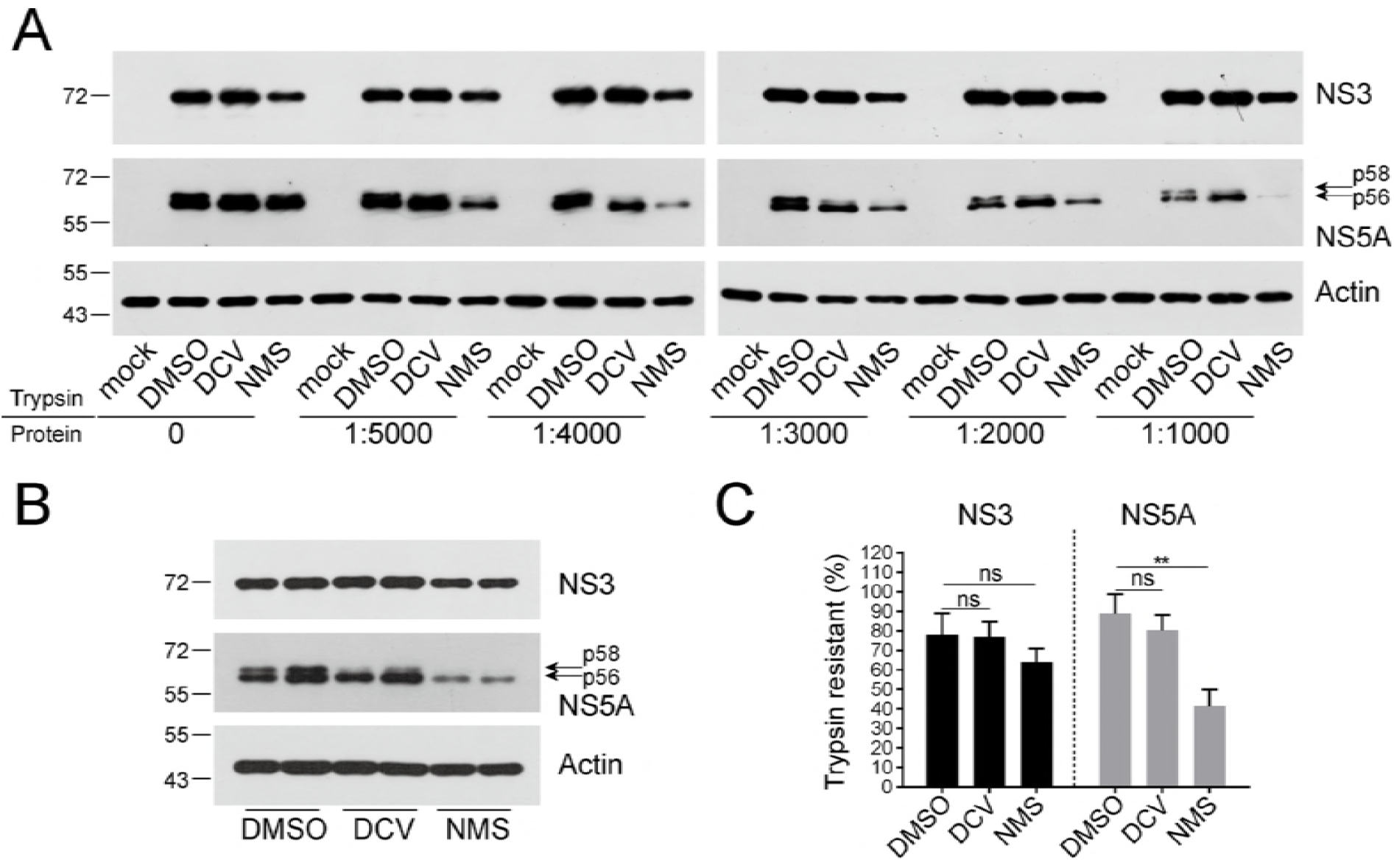
DCV does not change the tertiary structure of NS5A protein. (A) The sgJFH1 replicon cells were treated with DCV (10 nM) or NMS-873 (5 μM) for 4h. The cells were solubilized by 1% triton X-100 buffer and the cell lysates were incubated with trypsin at the indicated mass ratios (trypsin/total cellular protein; g/g) for 30min at 20°C. The digested cell lysates were analyzed by western blotting. Mock, Huh7.5 cells. (B) The digested cell lysates that were incubated with trypsin at a mass ration of 1:3000 (trypsin/total cellular protein) were analyzed by western blotting. (C)The basal phosphorylated form of NS5A (p56) and the protein level of NS3 in (B) were quantified. Mean values ± SD are shown (n=3). Statistical analysis was performed between the treated groups and the mock-treated groups (DMSO) (ns, not significant; **, *P*<0.01; two-tailed, unpaired *t*-test.). Similar results were observed in multiple independent experiments.

### Sensitivity of the replicase components to proteinase K (PK) digestion and the quaternary structure of the replicase

We propose that DCV may act on NS5A to affect the quaternary structure of the viral replicase. To monitor the quaternary structure change of the viral replicase, we used limited proteinase K (PK) digestion of the viral replicase. PK has broad specificity and a protein’s sensitivity to PK digestion reflects its accessibility to the enzyme. The change of a viral protein’s sensitivity to the PK digestion may reflect a positional change within a protein complex, and thus be used as an indicator of a quaternary structure change of a protein complex (30). To examine if the PK sensitivity reflects the quaternary structure change of the HCV replicase, we first compared the PK sensitivity of the replicase induced by expression of NS3-5B and the uncleaved NS3-5B polypeptide that is due to the inactivation of the NS3 (S139A) proteinase, as the unprocessed polyprotein may like YFV, can’t induce competent replicase assembly and sensitive to PK digestion (31). We first permeabilized the cells by digitonin at a concentration of 50 μg/ml. Under these conditions the viral replicase is preserved and active for viral replication (32). Then we digested the cells with proteinase K (PK) at a concentration (10 μg/ml) (Fig. 4A), under which, the C-terminal cytoplasmic domain of Calnexin could be efficiently digested while the N-terminal domain residing in the endoplasmic reticulum (ER) lumen remains intact (Fig. 4B). The sensitivity of NS3 or NS5A to the PK digestion was monitored by Western blotting with monoclonal antibodies against the NS3 helicase domain and the NS5A domain III (33), respectively. In the NS3-5B expressing cells, the majority of the NS5A was readily digested by PK. On the other hand, NS3 was much more resistant to PK digestion (Fig. 4B, left panel), although we detected NS3-specific bands with lower molecular weight than the full-length NS3 (Fig. 4B, asterisks). As the anti-NS3 antibody recognizes the helicase domain, these proteolytic fragments probably contain the helicase domain of NS3. In the cells expressing the HCV NS3-5B.S139A, similar to the YFV (31), the uncleaved HCV NS3-5B was readily digested by PK digestion (Fig. 4B).

**Figure 4.**
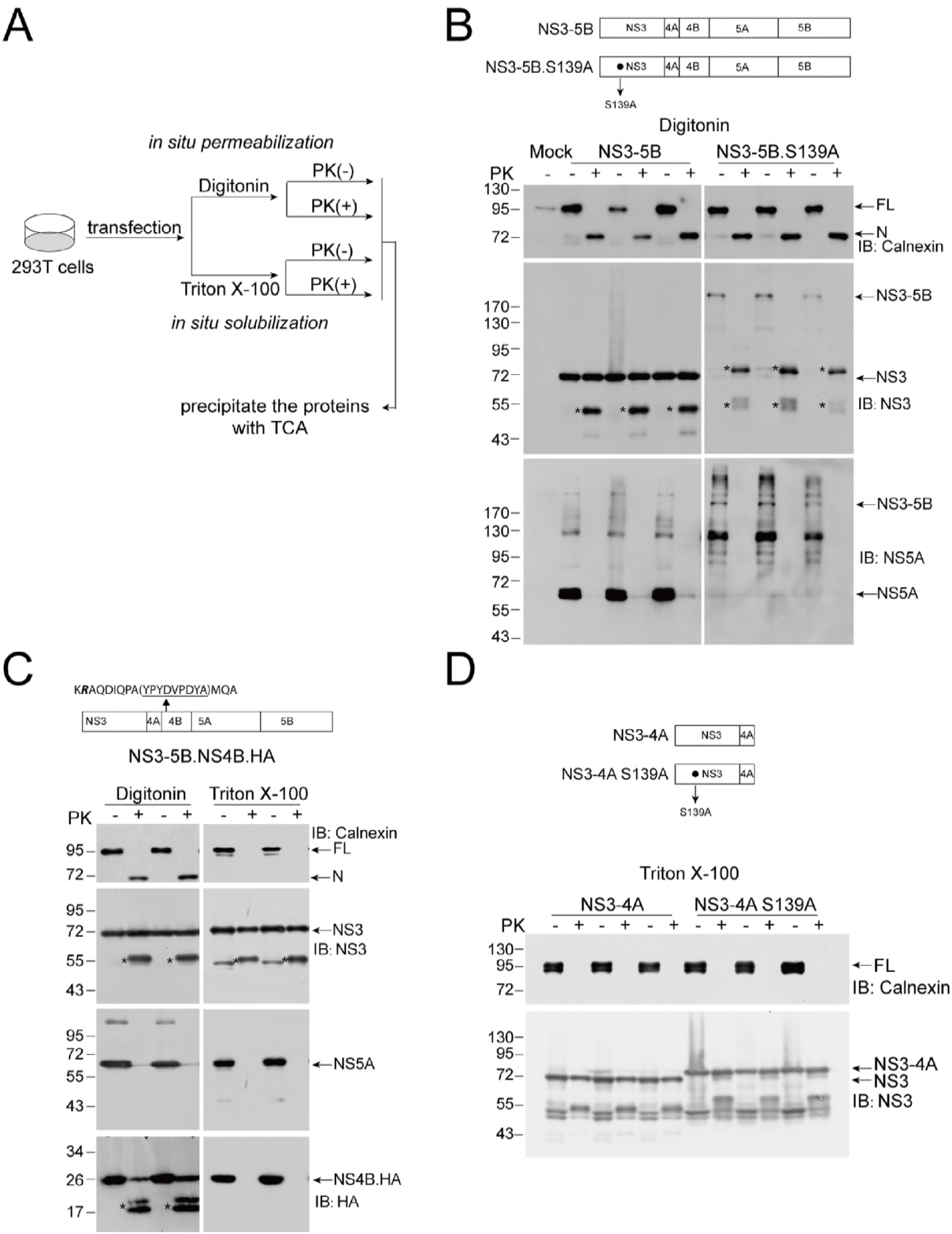
Sensitivity of replicase components to proteinase K (PK) digestion reflects the quaternary structural changes of the replicase. (A) The schematic of proteinase K digestion. (B) HEK293T cells expressing HCV NS3-5B or HCV NS3-5B.S139A mutant in triplicated wells were permeabilized with 50 μg/ml digitonin, scraped, and resuspended in buffer C. The permeabilized cells were treated (+) or not (-) with 10 μg/ml proteinase K (PK) for 5min at 37°C. Proteins were precipitated by TCA and analyzed by Western blotting. Mock, mock-transfected. Similar results were obtained in multiple independent experiments. (C) HEK293T cells expressing HCV NS3-5B.NS4B.HA in duplicated wells were permeabilized with 50 μg/ml digitonin or solubilized with 1% Triton X-100 in buffer C, scraped, and then treated (+) or not (-) with 10 μg/ml proteinase K (PK) for 5min at 37°C. Proteins were precipitated by TCA and analyzed by Western blotting. (D) HEK293T cells expressing HCV NS3-4A or NS3-4A.S139A mutant in triplicated wells were solubilized with 1% Triton X-100 in buffer C, scraped, and then treated (+) or not (-) with 10 μg/ml proteinase K (PK) for 5min at 37°C. Proteins were precipitated by TCA and analyzed by Western blotting. Representative data from multiple experiments with similar results are shown. The asterisks indicate proteolytic fragments. FL, full-length calnexin; N, protected N-terminal calnexin. The values to the left of the blots in panels B, C are molecular sizes in kilodaltons.

Considering the insensitivity of the viral replicase to PK may also be due to the enclosure of the viral proteins by the membranous replication complex (32), we also examined the sensitivity of the HCV NS5A, NS3 and NS4B to PK when the cells were treated by 1% Triton X-100 to disrupt the intracellular membranes. To facilitate the detection, we expressed the NS3-5B plasmid containing an HA-tagged NS4B (Fig. 4C), which was sub-cloned from a replication competent subgenomic replicon that contains the HA tag inserted between the AH1 and AH2 of NS4B (22, 34), ensuring the expression of NS3-5B.NS4B.HA is physiologically relevant. The cells expressing the NS3-5B.NS4B.HA were treated with digitonin and Triton X-100 respectively, and then digested by PK. The viral proteins’ sensitivity to PK were analyzed by Western blotting with antibodies against NS3, NS5A and HA (for detection of NS4B). We found that in the Triton X-100-treated cells, Calnexin was completely digested by PK, indicating the disruption of the ER lumen (Fig. 4C). Most NS3 were resistant to PK digestion in a similar way as in the digitonin-treated cells (Fig. 4C). A small amount of NS5A was resistant to PK digestion in the digitonin-treated cells whereas in the Triton X-100-treated cells, NS5A was completely digested by PK (Fig. 4C). Part of NS4B was resistant to PK digestion in the digitonin-treated cells whereas almost all the NS4B was digested by PK in the Triton X-100-treated cells (Fig. 4C). We also detected NS4B-specific proteolytic fragments in the digitonin-treated cells (Fig. 4C, bottom panel, asterisks), which may contain the N-terminal portion of NS4B, given the location of the HA tag (34). These data suggest that the resistance of NS3 to PK was not due to the membrane-mediated protection and the resistance of NS5A and NS4B to PK was membrane-dependent (see discussion).

As the NS3 expressed in the context of NS3-5B was resistant to PK in the TritonX-100-treated cells, we further examined if NS3 expressed alone also resistant to PK. We expressed HCV NS3-4A and the uncleaved NS3-4A.S139A in HEK293T cells. We solubilized the cells by 1% Triton X-100, and then treated the cells with PK as above. As in the NS3-5B expressing cells, both NS3 and the unprocessed NS3-4A were resistant to PK (Fig. 4D), suggesting the resistance of NS3 to PK digestion is probably due to its intrinsic folding (see discussion). Taken these data together, the sensitivity of the NS5A and NS4B to the PK digestion reflects the quaternary structure changes of the replicase.

### DCV induces quaternary structural changes of the viral replicase

We then examined the HCV viral proteins’ sensitivity to PK digestion in the presence of DCV (Fig. 5A). The cells expressing the NS3-5B were treated with 10 nM DCV upon transfection for 24 hours and then permeabilized, digested with PK as described above. DCV treatment didn’t significantly affect the sensitivity of NS3 and NS5A to PK digestion (Fig. 5B and 5C).

**Figure 5.**
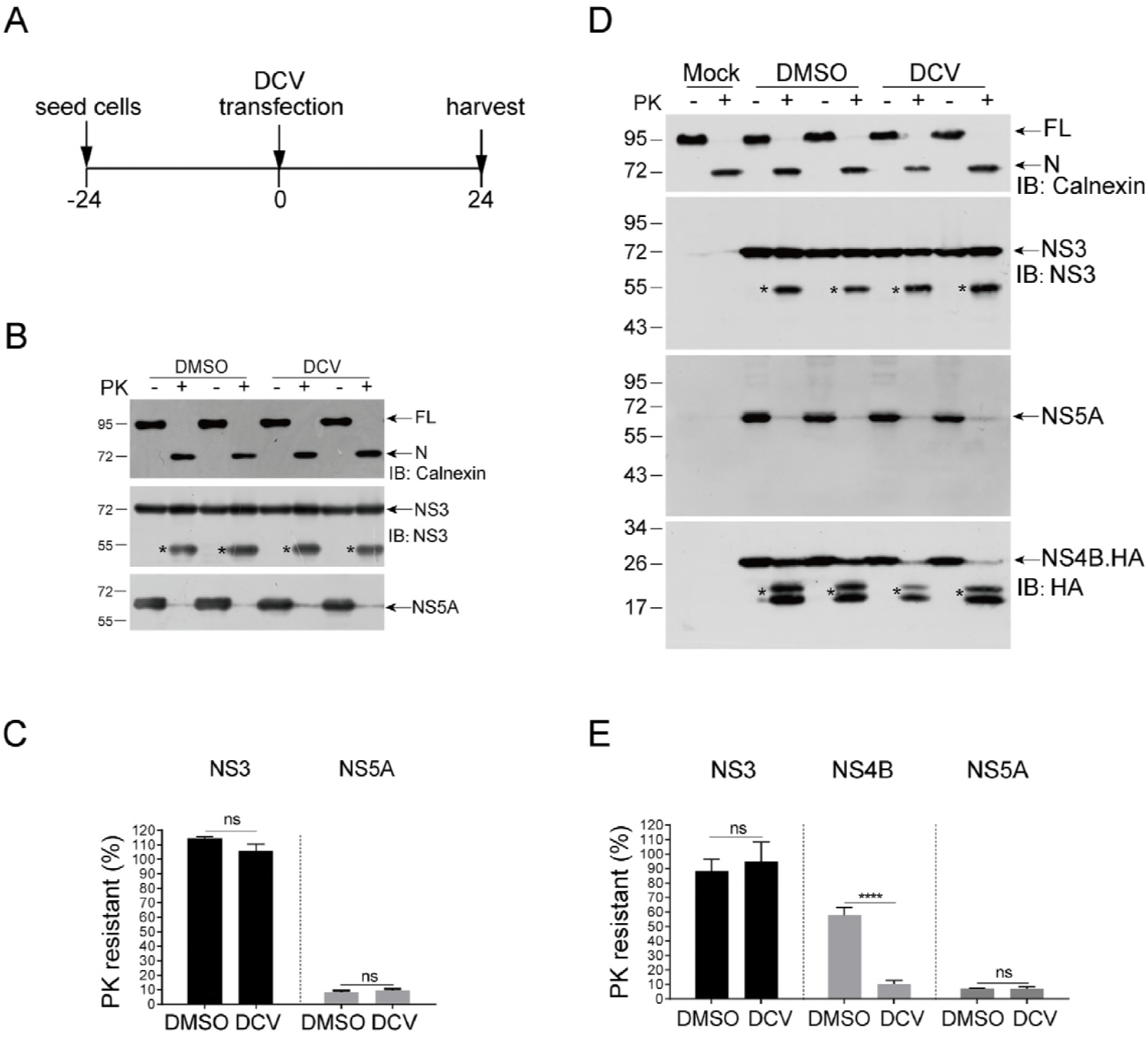
DCV induces quaternary structural changes of the viral replicase. (A) Experiment design for B and D. (B-C) HEK293T cells expressing HCV NS3-5B were treated with DCV (10 nM) for 24 hours. (D) The cells were permeabilized by 50 μg/ml digitonin and digested by proteinase K. The proteins were precipitated by TCA and analyzed by Western blotting. Representative data are shown. (C) The protein abundances were quantified and the PK-resistant efficiency was calculated as the ration of the undigested protein (PK+) to the total protein (PK-). Mean values ±SD are shown (n=3). (D-E) HEK293T cells expressing HCV NS3-5B.NS4B.HA were treated with DCV (10 nM) for 24 hours. Mock, mock-transfected. (D) Cells were permeabilized by digitonin and digested by proteinase K. The proteins were analyzed by Western blotting. Representative data are shown. (E) The protein abundances were quantified and the PK-resistant efficiency was calculated as the ration of the undigested protein (PK+) to the total protein (PK-). Mean values ±SEM are shown (n=6). Statistical analysis was performed between the treated groups and the mock-treated groups (DMSO) as indicated (ns, not significant; ****, *P*<0.0001; two-tailed, unpaired *t*-test.). The asterisks indicate proteolytic fragments. FL, full-length calnexin; N, protected N-terminal calnexin. The values to the left of the blots in panels B and D are molecular sizes in kilodaltons.

Then we sought to map the sensitivity of another viral replicase component, NS4B, to PK digestion in the presence of DCV. The HEK293T cells expressing the NS3-5B.NS4B.HA were treated with DCV, permeabilized and digested by PK. Consistent with what was seen when the wild type rather than HA tagged version of NS3-5B was expressed, DCV treatment didn’t significantly change the sensitivity of NS3 and NS5A to PK digestion (Fig. 5D and 5E). Strikingly, DCV treatment significantly increased the sensitivity of NS4B to PK digestion (Fig. 5D and 5E), which is probably due to quaternary structural changes of the viral replicase and exposure of NS4B.

### DCV impairs NS4B-involved protein interactions within the viral replicase

Given that DCV induced quaternary structural changes of the viral replicase, we hypothesized that the quaternary structure changes of the viral replicase may be coupled with changes in the protein-protein interactions within the viral replicase components. We used immunoprecipitation (IP) to examine the protein-protein interactions among the replicase components using both subgenomic replicon and NS3-5B expressing systems that contain HA-tagged replicase components. We first treated the sgJFH1.NS5A.HA replicon cells (22) (Fig. 6A) with DCV (10 nM) and NMS-873 (5 μM) for 4 hours and then solubilized the cells and captured the HA-NS5A. The NS3 protein was efficiently co-immunoprecipitated and the relative efficiency of co-precipitation with NS5A in the DCV- and NMS-873-treated samples were comparable with that in the DMSO-treated samples (Fig. 6A), suggesting that DCV and NMS-873 do not affect the NS3-NS5A interaction in replicon cells. As DCV affects the early biogenesis of the replicase (19) but not the pre-assembled replicase (15), we then added the DCV before replicase assembly in the replicase assembly surrogate system that expresses the NS3-5B.NS5A.HA, and then examined the NS3-NS5A interaction. In this system, in the DMSO-treated cells, NS3 was co-immunoprecipitated with HA-NS5A at a similar IP efficiency (about 30%) as in the replicon cells (Fig. 6A and 6B). DCV and NMS-873, as in the replicon cells, didn’t significantly change the NS3-NS5A interaction (Fig. 6B).

**Figure 6.**
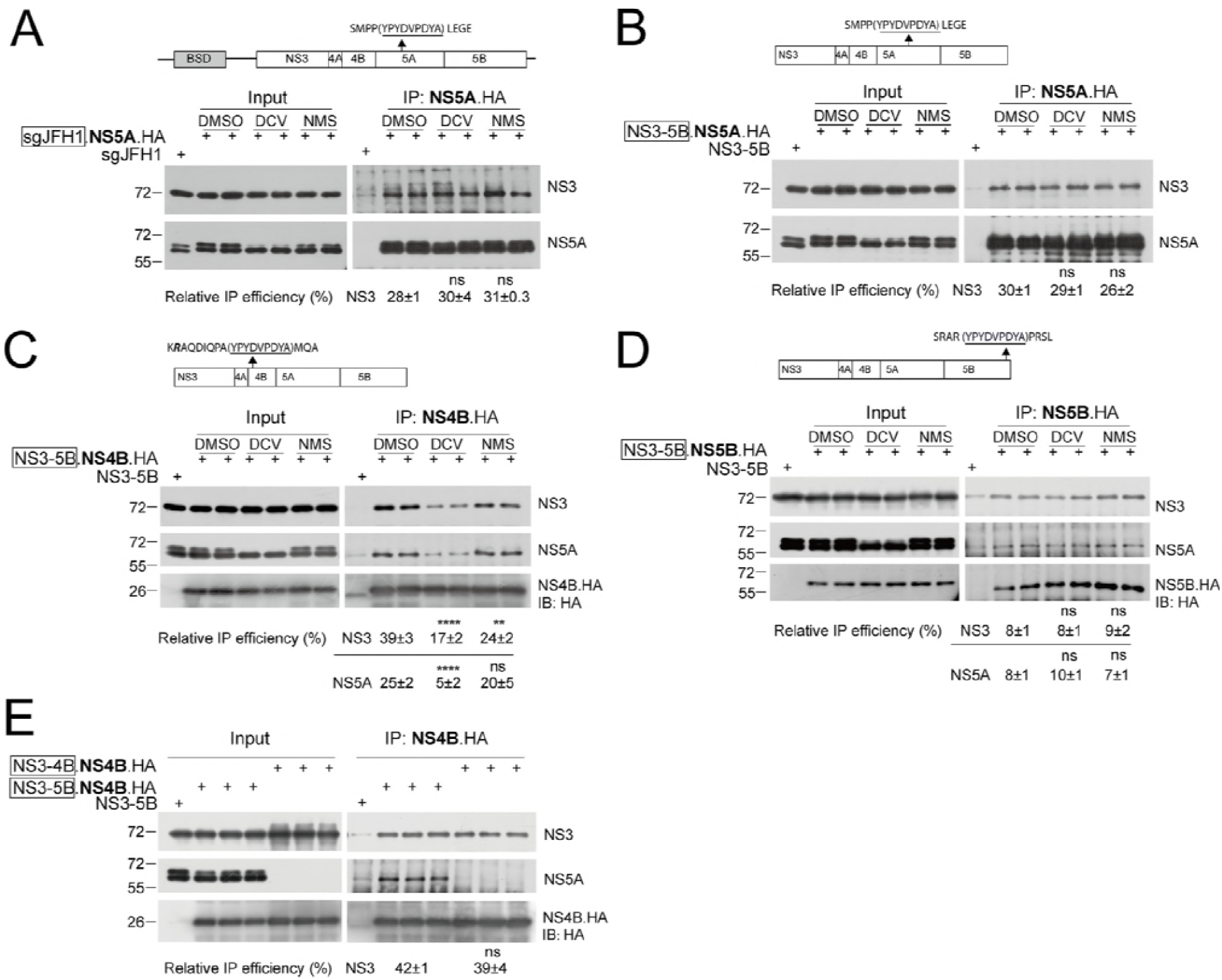
DCV impairs NS4B-NS5A and NS4B-NS3 interactions. (A) The sgJFH1-NS5A.HA cells were treated with DCV (10 nM) or NMS-873 (5μM) for 4h. The solubilized cell lysates were incubated with anti-HA beads. The captured proteins were analyzed by Western blotting with the antibodies indicated. The cell lysate from sgJFH1 was used as a negative control. The protein abundances were quantified. The relative immunoprecipitation (IP) efficiency was calculated as (immunoprecipitated NS3/ input NS3)/(captured NS5A/input NS5A). Mean values ± SD are shown (n=2). (B) HEK293T cells expressing HCV NS3-5B.NS5A.HA were treated with DCV (10 nM) for 24 hours or NMS-873 (5 μM) for 4 hours. The DCV was added upon transfection. The cell lysates were captured by anti-HA beads as described in (A). The cell lysate from 293T transfected with plasmids expressing HCV NS3-5B was used as a negative control. Representative picture are shown. The protein abundances were analyzed and the relative IP efficiency was calculated as in (A). Mean values ± SEM are shown (n=4). (ns, not significant; two-tailed, unpaired *t*-test.). (C) HEK293T cells expressing HCV NS3-5B.NS4B.HA were treated with DCV (10 nM) for 24 hours or NMS-873 (5 μM) for 4 hours as above. Representative picture of multiple experiments are shown. The relative immunoprecipitation (IP) efficiency was calculated as (immunoprecipitated NS3 or NS5A/ input NS3 or NS5A)/ (captured NS4B.HA /input NS4B.HA). Mean values ± SEM are shown (n=8). (D) HEK293T cells expressing HCV NS3-5B.NS5B.HA were treated with DCV (10 nM) for 24 hours or NMS-873 (5 μM) for 4 hours as above. Representative picture of multiple experiments are shown. The relative immunoprecipitation (IP) efficiency was calculated as (immunoprecipitated NS3 or NS5A/ input NS3 or NS5A)/(captured NS5B.HA /input NS5B.HA). Mean values ± SEM are shown (n=6). Statistical analysis was performed between the treated groups and the mock-treated groups (DMSO) as indicated (ns, not significant; **, *P*<0.01; ****, *P*<0.0001; two-tailed, unpaired *t*-test.). (E) HEK293T cells were transfected with plasmids expressing HCV NS3-5B.NS4B.HA or NS3-4B.NS4B.HA. The cell lysates were captured by anti-HA beads and the captured proteins were analyzed by Western blotting. The relative immunoprecipitation (IP) efficiency was calculated as (immunoprecipitated NS3 or NS5A/ input NS3 or NS5A)/ (captured NS4B.HA /input NS4B.HA). Mean values ±SD are shown (n=3). Statistical analysis was performed between the NS3-4B.NS4B.HA groups and the NS3-5B.NS4B.HA groups. (ns, not significant; two-tailed, unpaired *t-*test.) The “input” panel and the “IP” panel were from a same film with different exposure time while the quantification was performed under the same exposure condition. The values to the left of the blots are molecular sizes in kilodaltons.

We then examined other protein-protein interactions within the viral replicase components. We used NS3-5B.NS4B.HA plasmid containing the HA-tagged NS4B and NS3-5B.NS5B.HA plasmid containing the HA-tagged NS5B, which was sub-cloned from a subgenomic replicon that contains the HA-tagged NS5B (22), ensuring that the expression of NS3-5B.NS5B.HA is physiologically relevant. We transfected the plasmids into HEK293T cells and captured the HA-tagged NS4B or NS5B, and then detected the co-immunoprecipitated NS5A and NS3. Strikingly, DCV significantly reduced the NS3-NS4B and NS4B-NS5A interactions, as evidenced by the reduced relative co-IP efficiencies of both NS3 and NS5A when NS4B was IP’d (Fig. 6C). NMS-873 reduced the interaction of NS3 with NS4B but not the interaction of NS5A with NS4B (Fig. 6C). In contrast to NS4B-mediated interactions, DCV and NMS-873 both didn’t significantly affect the interaction of NS3 or NS5A with NS5B (Fig. 6D). We noticed that the interactions of NS3 and NS5A with NS5B with were much less efficient than with NS4B (Fig. 6C and 6D).

Given that DCV targets NS5A yet impairs NS4B-NS3 interaction, we examined if NS5A affects the NS4B-NS3 interaction *per se*. We compared the NS4B-NS3 interaction in the context of NS3-4B and NS3-5B. The co-IP efficiencies of NS3 with NS4B were comparable in both situations (Fig. 6E), suggesting that the NS5A does not affect the NS4B-NS3 interaction *per se*. Taken together, these data suggest that DCV impairs NS4B-involved protein interactions amongst the viral replicase in the context of NS3-5B (see discussion).

### The NS5A Y93H variant is refractory to DCV-induced phenotypes and attenuates viral replication

To demonstrate the specificity of the DCV-induced phenotypes, we first tested NS4B’s sensitivity to PK digestion using the plasmid NS3-5B.NS4B.HA harboring the DCV-resistant mutation Y93H in NS5A both in HEK293T and Huh7 cells. Under DMSO treatment, the NS4B’s sensitivity to PK in the wild-type (WT) NS3-5B context and the context of NS3-5B.Y93H were comparable (Fig. 7A and 7B) in both HEK293T and Huh7 cells. DCV significantly affects NS4B’s sensitivity to PK in the wild-type (WT) NS3-5B context in both cell lines. In contrast, DCV didn’t affect NS4B’s sensitivity to PK in the context of NS3-5B.Y93H (Fig. 7A and 7B).

**Figure 7.**
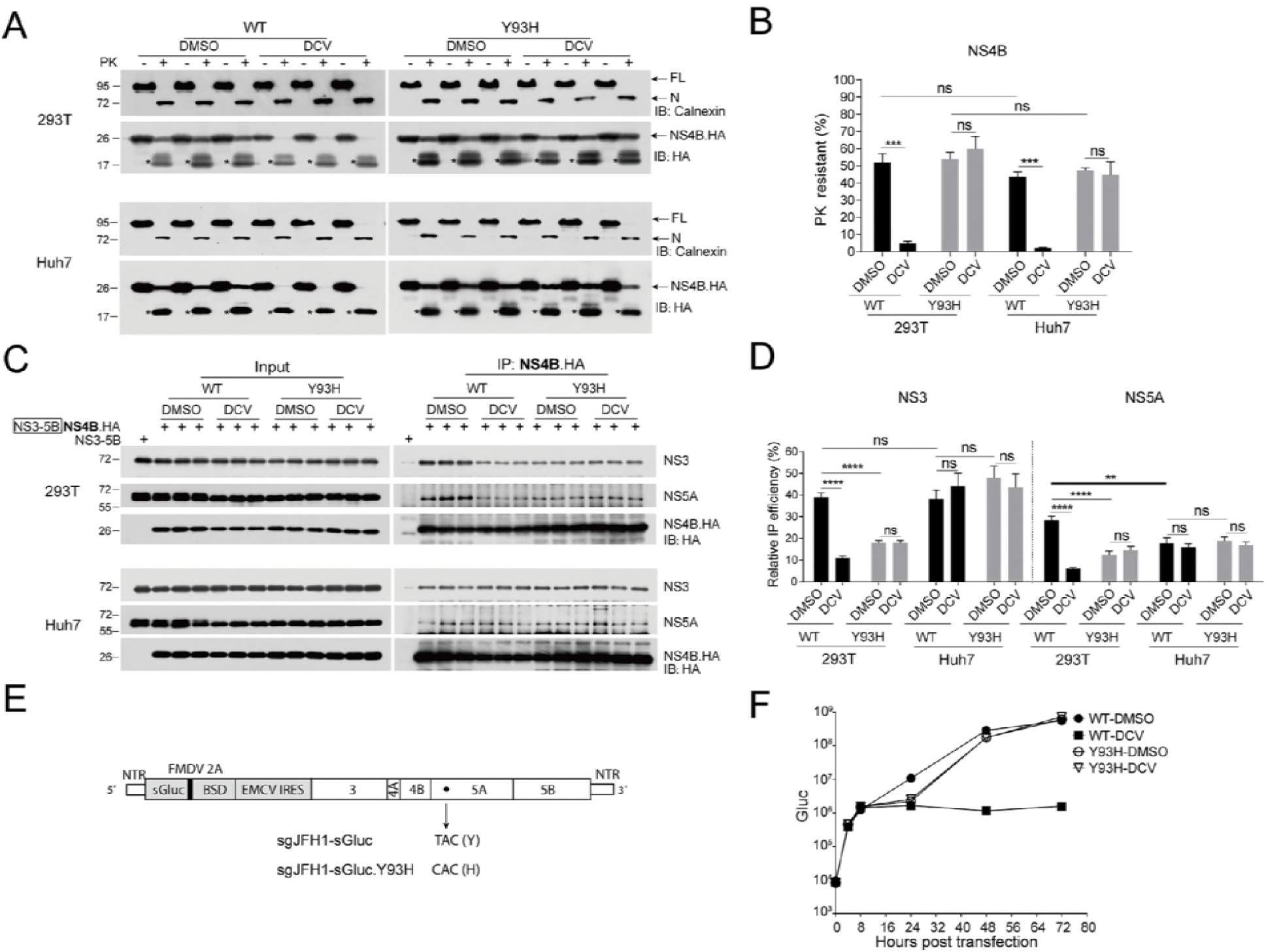
The NS5A Y93H variant was refractory to DCV-induced quaternary structural change of the viral replicase and changes of NS4B-mediated protein interactions within the replicase. (A, B) HEK293T and Huh 7 cells expressing HCV NS3-5B.NS4B.HA (WT) or NS3-5B.NS4B.HA Y93H were treated with DCV (10 nM) for 24 hours. Cells from triplicated wells were permeabilized with digitonin and treated with 10 μg/ml proteinase K (PK)(+) or no treatment (-). (A) Proteins were precipitated by TCA and analyzed by Western blotting. Representative data from multiple experiments with similar results are shown. The asterisk indicates NS4B-specific proteolytic fragments. FL, full-length calnexin; N, protected N-terminal calnexin. (B) The protein abundances in (A) were quantified and the PK-resistant efficiency was calculated as the ration of the undigested protein (PK+) to the total protein (PK-). Mean values ± SD are shown (n=3). (C, D) HEK293T and Huh7 cells expressing HCV NS3-5B.NS4B.HA (WT) or NS3-5B.NS4B.HA Y93H were treated with DCV (10 nM) for 24 hours. (C) The cell lysates were captured by anti-HA beads and the captured proteins were analyzed by Western blotting. Data from triplicated wells are shown. (D) The protein abundances in (C) were analyzed and the relative IP efficiency was calculated as described above. Mean values ± SEM are shown (n=6). Statistical analysis was performed between the treated groups and the mock-treated groups (DMSO) (ns, not significant; ***, *P*<0.001; ****, *P*<0.0001; two-tailed, unpaired *t*-test). The “input” panel and the “IP” panel were from a same film with different exposure time while the quantification was performed under the same exposure condition. (E) The schematic of sgJFH1-sGluc and sgJFH1-sGluc.Y93H. sGluc, *Gaussia* luciferase; NTR, non-translated region; BSD, balsticidin resistant gene; FMDV 2A (black bar). (F) Transient replication of sgJFH1-sGluc and sgJFH1-sGluc.Y93H. Huh7.5 cells were transfected with *in-vitro*-transcribed sgJFH1-sGluc or sgJFH1-sGluc.Y93H RNAs in the presence of DCV (10 nM). The luciferase activity in the supernatants was measured at the time points indicated. Mean values ± SEM are shown (n=4). The values to the left of the blots in panels A and C are molecular sizes in kilodaltons.

We then tested the impact of DCV on NS4B-involved interactions. In HEK293T cells, DCV reduced the NS4B-NS5A and NS4B-NS3 interactions in the context of wild-type (WT) NS3-5B but not in the context of NS3-5B containing the NS5A Y93H DCV resistance mutation (Fig. 7C and 7D). Whilst in Huh7 cells, DCV didn’t impair NS4B-NS5A and NS4B-NS3 interactions (Fig. 7C and 7D). It was noteworthy that under DMSO treatment, the IP efficiency of NS4B-NS5A in Huh7 cells was significantly lower than in the HEK293T cells (Fig. 7C and 7D) (see discussion). These data suggest the specificity of DCV’s effects on NS4B’s sensitivity to PK and the NS4B-involved protein-protein interactions in HEK293T cells.

We noticed that in the mock-treated cells (DMSO), the Y93H mutation reduced the interactions of NS4B with NS3 and NS5A in HEK293T cells (Fig. 7C, upper panel). We then examined if the Y93H mutation affects viral replication. We used a bi-cistronic subgenomic replicon of JFH1, expressing sGluc as a reporter (Fig. 7E). We compared the transient replication kinetics of WT and Y93H replicons in the presence or absence of DCV. As expected, the DCV only reduced the replication of WT replicon but not the Y93H replicon (Fig. 7F). In the mock-treated cells, compared with wild-type, the replication of the Y93H replicon was decreased about 5 fold at 24 hours post transfection, but caught up with the wild-type in the late time points (Fig. 7F), suggesting that the Y93H mutation attenuates the viral replication at early time points.

## Discussion

By imaging techniques, it has been demonstrated that DCV either changes the size of the NS5A-clustering sites (15) or not (19) and induces redistribution of NS5A to lipid droplets (17); DCV treatment alters HCV replication complex biogenesis, reduces the diameters of DMVs or completely blocks the DMV biogenesis when early treatment (19). In this study, we attempted to dissect the underlying mechanistic actions of DCV by biochemical experiments.

### DCV’s impact on the multimerization and folding of NS5A

NS5A dimerizes through the DI (8, 9). The dimeric NS5A may homodimerize to form oligomers or polymers in viral replicating cells, as the heterogeneous oligomeric state of the NS5A DI in solution has been observed (8). Pharmacological inhibitor studies suggest the existence of NS5A polymers within the cells (25). Although there is no effect of DCV on the NS5A dimerization (19), it is possible that DCV may affect the formation of NS5A polymers. We recently uncovered the aggregation-prone property of NS5A, and ablation of VCP, a member of the ATPases associated with diverse cellular activities (AAA+ ATPase family) results in aberrant aggregation of NS5A (26). Unlike the VCP inhibitors, DCV didn’t induce the aberrant aggregation of NS5A (Fig. 2). We mapped the structural change of NS5A by its sensitivity to limited trypsin digestion and found that DCV unlikely affect the overall structure of NS5A (Fig. 3). In contrast, the VCP inhibitor NMS-873 induces aggregation of NS5A and may cause an overall structural change to NS5A, resulting in increased sensitivity of NS5A to trypsin digestion (Fig. 2 and 3).

### PK sensitivity and quaternary structural changes of the replicase

We monitor the quaternary structural change of the viral replicase in a replicase assembly surrogate system by proteinase K (PK) digestion of the digitonin-permeabilized cells. The resistance of the replicase components to PK digestion may be either mediated by the membranous replication complex proposed previously (32) or by the proteinacous replicase (Fig. 8). As the HCV replicase induces protrusion of the ER toward the cytosolic side (4), the membrane-mediated PK resistance of the viral repilcase components should engage further membrane modification to form a DMV (4), which is probably mediated by enrichment of lipids such as cholesterol and PI4P (34, 35). While, when the HCV subgenomic replicon cells were permeabilized by diginotin as in this study, substantial replicative double stranded RNAs (dsRNA) resided within the DMVs became sensitive to RNase A and RNase III (36), indicating exposure of the replicase in the diginotin-permeabilized cells. The partially disruption of the DMVs may be due to the selective solubilization of the cholesterol by digitonin (37). Thus in the digitonin-permeabilized cells, the resistance of the replicase components to PK digestion may also partially be mediated by the proteinacous replicase (Fig. 8). The resistance of NS3-4A to PK digestion is membranes independent (Fig. 4D) and probably due to the unique folding of NS3-4A as the GFP protein (38). Only a small amount of NS5A was resistant to PK digestion (Fig. 4B and 4C), which may be explained by only a small amount of NS5A is engaged in the replicase assembly within the DMVs. The monoclonal antibody used for detection of NS5A recognizes the NS5A domain III (33). We can’t rule out the possibility that the largely unfolded NS5A domain III is at the outside of the replicase and more sensitive to PK and the domain I and domain II of NS5A are in a position within the replicase that renders them to be resistant to PK digestion. We tested this possibility by several commercial antibodies against the NS5A domain I, but unfortunately the antibodies didn’t work. About half of the NS4B was resistant to PK digestion (Fig. 4C, 5D and 5E) and the N-terminal NS4B-specific proteolytic fragments were detected (Fig. 4C, bottom panel, asterisks). We proposed that NS4B, especially the N-terminal portion of NS4B was protected together by the ER membrane and the ER-resident viral replicase (Fig. 8). In supporting this, DCV treatment selectively renders the full-length (mainly the C-terminal portion of) NS4B to be sensitive to PK while remains the N-terminal portion of NS4B (Fig. 5D, bottom panel, asterisks and Fig. 7A, asterisks) and NS5A unaffected (Fig. 5). We think these observations are unlikely due to the disruption of the DMV by DCV, as the disruption of DMV by 1% Triton X-100 makes the NS4B and NS5A were completely sensitive to PK (Fig. 4C). We proposed that DCV induces position change of the replicase relative to the ER membrane (see below), resulting in the exposure of the C-terminal portion of NS4B.

**Figure 8.**
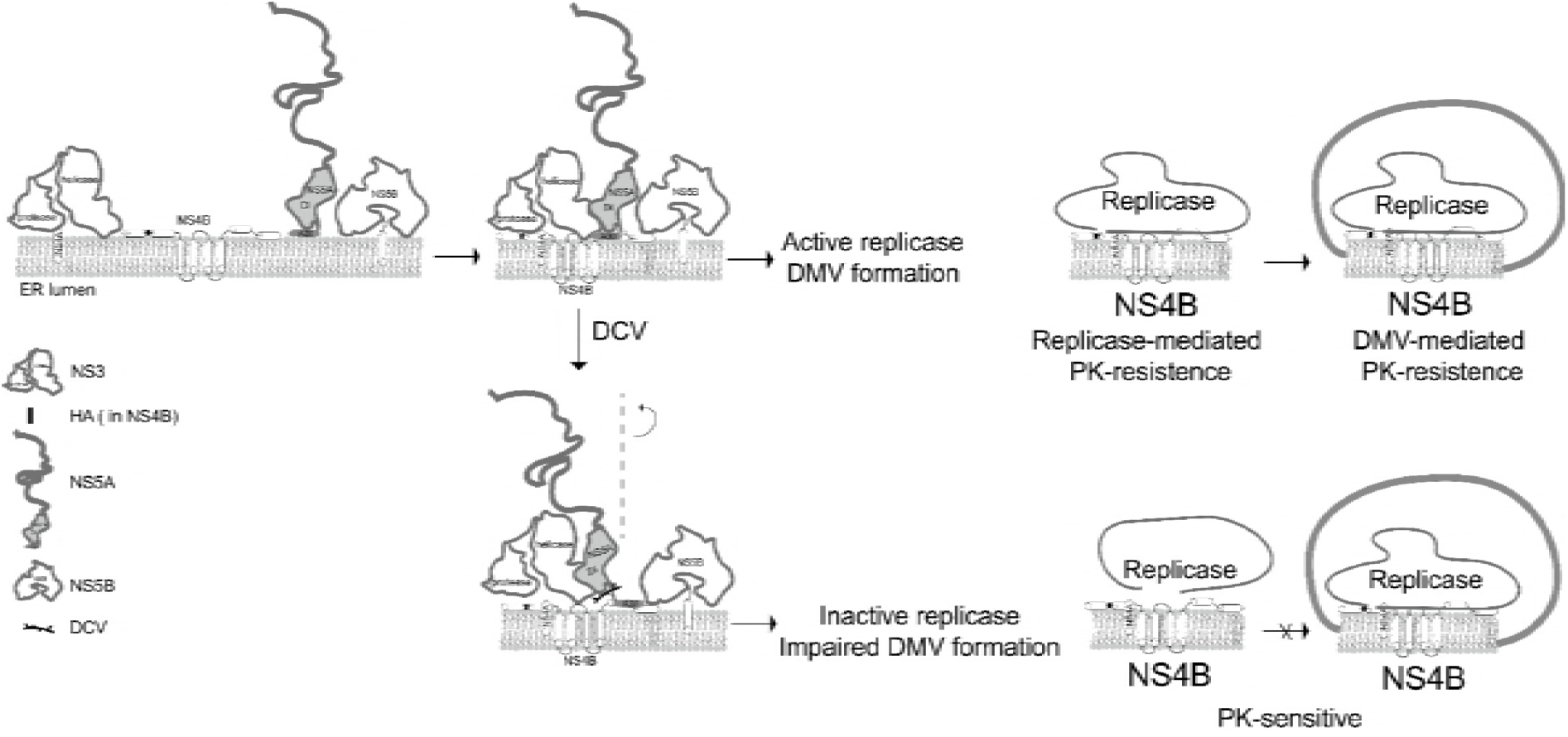
A proposed model of the DCV’s action on HCV replicase assembly. The viral proteins may exist as multimers *in vivo*. Only the monomers are shown. The NS3 consists of the N-terminal protease domain and the C-terminal helicase domain. NS3 associates with the membrane by interacting with NS4A. NS4B is a multi-spanning integral membrane protein. The HA tag (black bar) is inserted in-frame into between the first (AH1) and the second amphipathic helix (AH2). NS5A associates with membrane via the N-terminal amphipathic helix A30 (Grey). Concerted protein-protein interactions within the replicase components induce assembly of the active replicase on the endoplasmic reticulum (ER), resulting in protruding of the ER membrane. The protruded ER membrane is further modified to form a double membranous vesicle (DMV), namely the replication complex. DCV binding may cause a position change of NS5A relatively to the membrane. The position change of NS5A may be allosterially transmitted to NS3 via NS5A-NS3 interaction, resulting in the separation of the cytosolic domains of NS5A and NS3 from the membrane-resident NS4B and reduction of the NS4B-NS3 and NS4B-NS5A interactions. Alternatively, the position change of NS5A may be allosterically transmitted to NS4B via NS5A-NS4B interaction, resulting in a conformation change of NS4B and reductions of NS4B-involved protein-protein interactions. Aberrant protein-protein interactions within the replicase components fail to assemble active replicase. The resistance of NS4B to proteinase K (PK) digestion may be either mediated by the proteinaceous replicase or by the double-membranous replication complex.

### DCV’s effect on the protein-protein interactions within the replicase and discrepancy of the effect on the NS4B-NS5A interaction in different cellular contexts

Mechanistically, DCV impaired the NS4B-involved protein-protein interactions (NS4B-NS3 and NS4B-NS5A) within the replicase but not the NS5A-NS3, NS5B-NS5A and NS5B-NS3 (Fig. 6). Given that NS5A doesn’t affect NS4B-NS3 interaction *per se* (Fig. 6E), DCV may allosterically affect NS4B-NS3 interaction (see below). It should be noted that we only observed the DCV’s effect on the NS4B-involved protein-protein interactions in HEK293T cells but not in Huh7 cells (Fig. 7C and 7D). These might be due to the differences of the lipid metabolism and different lipid environments of these two cell lines as the lipids play crucial roles in the viral proteins and virus-host interactions for the replicase assembly (35). HEK293T cells support replication of JFH1 subgenomic replicons (24), making it physiologically relevant to study the replicase assembly in these cells. We noticed that the protein-protein interactions within the replicase components are comparable without significant differences except for the NS5A-NS4B interactions in HEK293T cells and Huh7 cells (Fig. 6A, 6B, 7B and 7D). There was less efficient NS5A-NS4B interaction in the Huh7 cells than in the HEK293T cells (Fig. 7D). The stronger NS5A-NS4B interaction in the HEK293T cells may make the DCV-induced allosteric change of NS4B visible in this study. We can’t rule out the existence of the allosteric change of NS4B in the Huh7 cells, which is probably invisible by our co-IP experiments. More sensitive methods should be employed to address this issue in the future studies.

### DCV’s action model

It has been proposed that DCV binds to the membrane-proximal side of NS5A (19) and this binding may perturb the positioning or orientation of NS5A to impair its interactions with other replicase components or host factors (19, 20, 39, 40). Our observations support this hypothesis. The quaternary structure change may be initiated by the insertion of the DCV into the membrane-proximal side of the dimeric NS5A. DCV binding may cause a position change of NS5A relatively to the membrane, separating the cytosolic domains of NS5A from membrane. Given that NS5A doesn’t affect NS4B-NS3 interaction *per se* (Fig. 6E), while in the context of NS3-5B, the position change of NS5A may be allosterically transmitted to NS3 via NS3-NS5A interaction, as NS3-NS5A interaction was not affected by DCV (Fig. 6A and 6B), separating NS3 and NS5A from the membrane-resident NS4B and resulting in the reduction of NS4B-NS3 and NS4B-NS5A interaction (Fig. 6C and 8). Alternatively, the position change of NS5A may be allosterically transmitted to NS4B via NS5A-NS4B interaction (41), resulting in a conformation change of NS4B and reductions of NS4B-involved protein-protein interactions (Fig. 8).

### Mechanism of DCV-resistant mutations

The DCV-resistance of Y93H mutant is partly explained by the reduction of binding to DCV (19). In this study, we found that Y93H mutation reduced NS4B-NS3 and NS4B-NS5A interactions (Fig. 6C, up panel), mimicking the DCV. The DCV-binding pocket might be formed by the ER membrane and the membrane-proximal side of NS5A. In the back-to-back dimeric NS5A, Y93 locates in the membrane-proximal region (19). It is possible that the Y93H mutation might somehow affect the position of NS5A and impair the binding cleft for DCV as well as attenuate the NS4B-NS3 and NS4B-NS5A interactions. The impaired NS4B-NS3 and NS4B-NS5A interactions may account for the reduced viral replication kinetics of the Y93H-containing replicon in the early time points (Fig. 7F).

In conclusion, we demonstrate that DCV induces a quaternary structural change of the HCV replicase, and allosterically impairs the protein-protein interactions among the viral replicase in a repilcase assembly surrogate system. As a highly conserved replication strategy is used by almost all of the positive-stranded RNA viruses, the DCV action mechanism elucidated here might enlighten the development of DAAs for other positive-stranded RNA viruses.

## Materials and Methods

### Plasmids

The plasmids phCMV-NS3-5B (2a) expressing the HCV NS3-5B (JFH1) polypeptide was reported previously (22). To generate the plasmid phCMV-NS3-5B.5A.HA expressing an HA-tagged NS5A in the context of NS3-5B, an SanDI/BsrGI digested fragment was swapped from the plasmid sgJFH1-NS5A.HA (22) into the similarly digested plasmid phCMV-NS3-5B. To generate the phCMV-NS3-5B.5B.HA expressing an HA-tagged NS5B in the context of NS3-5B, a fragment encompassing the BsrGI site to the end of the NS5B was amplified from the plasmid sgJFH1-DU-NS5B.HA1 (22) and then digested by BsrGI/EcoRI and ligated into the similarly digested plasmid phCMV-NS3-5B. To generate the phCMV-NS3-5B.4B.HA expressing an HA-tagged NS4B in the context of NS3-5B, a PmlI digested fragment was swapped from the plasmid sgJFH1-NS4B.HA into the similarly digested plasmid phCMV-NS3-5B. The DCV resistant mutation Y93H was introduced into the plasmid phCMV-NS3-5B.4B.HA by fusing PCR-mediated mutagenesis to get the plasmid phCMV-NS3-5B.4B.HA.Y93H. The plasmid phCMV-NS3-5A.4B.HA and the plasmid phCMV-NS3-4B.4B.HA were generated by PCR amplification of the NS3-5A region and the NS3-4B region from the plasmid phCMV-NS3-5B.4B.HA and then ligated into the BglII/EcoRI sites in the plasmid phCMV, respectively. The plasmid phCMV-NS3-4A were generated by PCR amplification of the NS3-4A region and then ligated into the BglII/EcoRI sites in the plasmid phCMV. The NS3 protease inactive mutation S139A was introduced into the plasmid phCMV-NS3-5B to get the plasmid phCMV-NS3-5B.S139A and phCMV-NS3-4A.S139A. HCV con1b subgenomic plasmid BB7 was reported previously (23). To generate the subgenomic replicon sgJFH1-sGluc expressing a secreted *Gaussia* luciferase (sGluc), using the sgJFH1 (22) as a backbone, first a cassette containing the sGluc-2A fragment was assembled with a cassette containing the BSD-EMCV IRES by fusing PCR and then the assembled fragments was digested by AgeI/KpnI and ligated into the similarly digested sgJFH1 to get the plasmid sgJFH1-sGluc. The Y93H mutation was introduced into the plasmid sgJFH1-sGluc by fusing-PCR mediated mutagenesis as described above. All of the constructs were proofed by DNA sequencing. Detailed information is available upon request.

### Cells

The human embryonic kidney cell line HEK-293T and human hepatoma cell line Huh 7(Cell Bank of the Chinese Academy of Sciences, Shanghai, China, www.cellbank.org.cn) were routinely maintained in Dulbecco’s modified medium supplemented with 10% FBS (Gibco) and 25mM HEPES (Gibco). Huh7.5 (provided by Charles Rice) was routinely maintained in a similar medium supplemented with non-essential amino acids (Gibco). The replicon cells were generated as reported previously (22), and routinely maintained with the medium supplemented with 0.5 μg/ml blasticidin.

### Virus

The HCVcc was generated as described previously (42). Briefly XbaI-linearized plasmids were purified and used as templates for the *in-vitro* transcription by MEGAscript T7 Transcription Kit (Ambion) according to the manufacturer’s protocol. The *in-vitro-*transcribed RNAs were electroplated into Huh7.5 cells. The viruses in the supernatants were collected and the virus titer was determined in Huh7.5 cells by limiting dilution.

### Antibodies and inhibitors

Anti-NS5A monoclonal antibody (9E10; gifted by Charlie Rice) which recognizes the domain III of NS5A (33) was used in Western and immunofluorescence analyses at 1:2,000 and 1:200 dilutions, respectively; Anti-NS3 monoclonal antibody (Virogen; 217-A) which recognizes the helicase domain (a.a. 1350-1460) was used at 1:1,000 dilution; Anti-β-actin antibody (Sigma; A1978) was used at 1:4,000 dilution; Anti-calnexin antibody (BD; 610523) was used at 1:2,000 dilution; Anti-HA antibody (Abcam; ab130275) was used at 1:000 dilution; Goat-anti-mouse HRP IgG (Santa Cruz; sc-2004) was used at 1:2,000 dilution; Goat-anti-rat HRP IgG (Santa Cruz; sc-2005) was used at 1:2,000 dilution. Alexa Fluor 488 goat-anti-mouse IgG (Life technologies; 1298479) was used at 1:200 dilution.

Daclatasvir (BMS-790052) was purchased from Selleckchem (S1482), NMS-873 was purchased from Selleckchem (S7285).

### Luciferase activity

Supernatants were taken from cell medium and mixed with equal volume of 2×; passive lysis buffer (Promega). Luciferase activity was measured with Renilla luciferase substrate (Promega) according to the manufacturer’s protocol.

### Transfection

For plasmid transfection, HEK 293T cells were seeding onto poly-L-lysine (Sigma)-coated 6-well plates at a density of 4.5E5 cells ml^−1^ and then transfected with plasmids using a TransIT-LT1 transfection kit (Mirus) according to the manufacturer**’**s protocol. For RNA transfection, Huh 7.5 cells were seeding onto 48-well plates at a density of 2.5E5 cells ml^−1^ and then transfected with 0.5μg *in-vitro*-transcribed RNA using a TransIT-mRNA transfection kit (Mirus) according to the manufacturer**’**s protocol.

### Western blotting

After washing with PBS, cells were lysed with 2×;SDS loading buffer (100 mM Tris-Cl [pH 6.8], 4% SDS, 0.2% bromophenol blue, 20% glycerol, 10% 2-mercaptoethanol) and then boiled for 5 min. Proteins were separated by SDS-PAGE and transferred to a nitrocellulose membrane. The membranes were incubated with blocking buffer (PBS, 5% milk, 0.05 % Tween) for 1 hour and then with primary antibody diluted in the blocking buffer. After three washes with PBST (PBS, 0.05 % Tween), the membranes were incubated with secondary antibody. After three washes with PBST, the membrane was visualized by Western Lightning Plus-ECL substrate (PerkinElmer, NEL10500). The protein bands were quantified by densitometry with Image J if necessary.

### Electron microscopy

Cells were fixed in 0.1M phosphate buffer containing 2.5 % glutaraldehyde for 2 hr, scrapped in phosphate buffer and then the cell pellets were washed with phosphate buffer. After postfixation with 1% osmium tetroxide in phosphate buffer at 4°C for 2 hr, the samples were dehydrated and embedded in Epoxy resins. Ultrathin sections were prepared using a Reichert ultramicrotome, contrasted with uranyl acetate and lead citrate, examined under a tecnai sprit transmission electron microscope (FEI) electron microscope at 60 KV.

### Semi-denaturing detergent agarose gel electrophoresis (SDD-AGE)

SDD-AGE was carried out essentially as described (26). Briefly, cells were lysed with lysis buffer (50 mM Hepes [pH 7.5], 150 mM NaCl, 2.5 mM EDTA, 1 % Triton X-100) containing 1×;Complete Protease Inhibitor (Roche). Then the cell lysates were passed through a 27-gauge needle 10 times and centrifuged at 3000 ×;*g* for 10min. The supernatant were mixed with 4×; loading buffer (2×;TAE, 20 % glycerol, 8 % SDS, bromophenol blue). The samples were separated by 1.5% agarose gel (1×;TAE with 0.1 % SDS) in running buffer (1×;TAE with 0.1 % SDS) on ice for 75min with a constant voltage of 50 V. The proteins were analyzed by Western blotting. The NS5A aggregation was analyzed essentially as described (26).

### Limited trypsin proteolysis

Cells in 6-well plates were lysed with 150ul lysis buffer (50 mM TrisCl [pH 7.5], 1 mM EDTA, 15 mM MgCl2, 10 mM KCl, 1% Triton X-100, proteinase inhibitor [Roche]). Cell lysates were passed through a 27-gauge needle 20 times and centrifuged at 12000 ×;*g* for 10min. The protein concentration of the cell lysates was quantified by Pierce BCA Protein Assay Kit (Thermo Scientific). The cell lysates were incubated with trypsin (Sigma) at various mass ratios (g/g) for 30 min at 20°C. The reaction was terminated by adding equal volume of 2×;SDS loading buffer. The samples were boiled and analyzed by Western blotting.

### Proteinase K digestion

Cells in 6-well plates were permeabilized with 1ml 50 μg/ml digitonin in buffer C (20 mM HEPES-KOH [pH 7.7], 110 mM potassium acetate, 2 mM magnesium acetate, 1 mM EDTA) at room temperature for 5min, then scraped into 400 μl buffer C; or cells were solubilized with 400 μl 1% Triton X-100 in buffer C at room temperature for 5min, and then incubated with or without 10 μg/ml proteinase K for 5min at 37°C. Proteins were precipitated by adding equal volume of 40% TCA (trichloroacetic acid). After centrifugation at 12000 ×;*g* for 10min, the precipitates were washed by 500 μl acetone, and then dissolved in 30 μl 2D buffer (7M Urea, 2M thiourea). The samples were mixed with equal volumes 2×;SDS loading buffer and boiled for 5 min and then analyzed by Western blotting.

### Immunoprecipitation

Cells in 6-well plates were lysed with 150 μl lysis buffer (50 mM TrisCl [pH 7.5], 1 mM EDTA, 15 mM MgCl2, 10 mM KCl, 1% Triton X-100, proteinase inhibitor [Roche]). Then cell lysates were passed through a 27-gauge needle 20 times and centrifuged at 12000 ×;*g* for 10min. 15 μl of the supernatant was taken and mixed with equal volume of 2×;SDS loading buffer as input (10%). The rest of the clarified cell lysates were incubated with 10 μ □anti-HA magnetic beads (Pierce, SB246262) overnight with rotation at 4°C. After four washes with wash buffer (50 mM TrisCl [pH 7.5], 1 mM EDTA, 15 mM MgCl2, 10 mM KCl, 1% Triton X-100), the beads were lysed with 2×;SDS loading buffer. The samples were boiled for 10 min then analyzed by Western blotting.

## Acknowledgements

We are grateful to Charles Rice (The Rockefeller University) for providing the HCV reagents; to Margaret R. MacDonald and Mohsan Saeed (The Rockefeller University) for their critical reading of the manuscript; to Wuhiu Song (Fudan university) for her excellent technical assistance.

